# Genetic predictors of gene expression associated with risk of bipolar disorder

**DOI:** 10.1101/043752

**Authors:** Kaanan P. Shah, Heather E. Wheeler, Eric R. Gamazon, Dan L. Nicolae, Nancy J. Cox, Hae Kyung Im

**Affiliations:** Section of Genetic Medicine, Department of Medicine, University of Chicago, Chicago IL, USA; Departments of Biology and Computer Science, Loyola University Chicago, Chicago IL, USA; Academic Medical Center, University of Amsterdam, Amsterdam, The Netherlands; Division of Genetic Medicine, Vanderbilt University, Nashville, TN, USA; Department of Statistics, University of Chicago, Chicago, IL, USA

## Abstract

Bipolar disorder (BD) affects the quality of life of approximately 1% of the population and represents a major public health concern. It is known to be highly heritable but large-scale genome-wide association studies (GWAS) have discovered only a handful of markers associated with the disease. Furthermore, the biological mechanisms underlying these markers need to be elucidated. We recently published a gene-level association test, PrediXcan that integrates transcriptome regulation data to characterize the function of these markers in a tissue specific manner. In this study, we developed prediction models for mRNA levels in 10 brain regions using data from the GTEx project and performed PrediXcan analysis in WTCCC as well as in an independent cohort, GAIN. We replicate the association between predicted expression of *PTPRE* and BD risk in whole blood and recapitulate the association in brain tissues. *PTPRE* encodes the protein tyrosine phosphatase, receptor type E, that is known to be involved in RAS signaling and activation of voltage-gated K+ channels. We also found a new genome-wide significant association between lower predicted expression of *BBX* (bobby sox homolog) in the anterior cingulate cortex region of the brain and increased risk of BD (p_WTCCC_ = 7.02 x 10^−6^, p_GAIN_ = 4.68 x 10^−3,^ p_meta_ = 1.11 x 10^−7^). In sum, we used our mechanistically informed approach, PrediXcan, to identify and replicate two novel genome-wide significant genes using existing GWAS studies.

## Introduction

Bipolar disorder (BD) is a psychiatric disease that severely affects the quality of life of approximately 1% of the population [1,2]. BD is characterized by episodes of mania and depression with onset in late adolescence. About 10-20% of individuals with BD commit suicide and nearly a third attempt suicide [3]. It is therefore clear that further understanding of the etiology of BD is necessary to address this major public health concern. Genetic studies to examine the biological mechanisms of the disease and its risk factors can generate knowledge that aid in improving diagnosis, treatment, and monitoring of individuals at risk for developing this disease.

Previous twin and family studies have shown that risk for BD is highly heritable with estimates of genetic factors explaining ~80% of disease risk [1,4]. Despite the clear genetic component to risk of BD, identifying specific genes and variants is challenging. Smaller scale GWAS studies (n < 5,000) had limited power and discovered only a handful of genome-wide significant loci, namely *DGKH* [5], *ANK3* [6], *CACNA1C* [6], and *NCAN* [7]. Subsequently, the Psychiatric Genetics Consortium (PGC) consolidated these studies to perform a large GWAS and meta-analysis for BD which included ~16,000 individuals in stage 1, and an additional ~64,000 individuals for replication. The consortium replicated the association near *CACNA1C* and identified a novel locus near *ODZ4* [8]. The most recent meta-analysis, including the PGC samples and an additional ~7,000 samples further identified 3 new loci near *ADCY2, POU3F2/MIR2113*, and *TRANK1* [9]. Thus, despite concerted efforts and the combination of data on ~24,000 individuals, there are still fewer than a dozen know BD risk loci.

As we begin to discover loci associated with disease risk, it is critical to understand the underlying mechanisms if we are to translate this knowledge to improve treatment. One such mechanism is through altered gene expression. Numerous studies have identified single nucleotide polymorphisms (SNPs) that are associated with variation in gene expression levels across a variety of tissues [10-12]. These so-called expression quantitative trait loci (eQTLs) are enriched among GWAS-identified loci across diseases [13], and specifically in BD [14]. Thus, there is growing consensus that the genetic regulation of gene expression plays an important role in complex disease risk.

An alternative approach to GWAS for dissecting disease risk is to identify sets of differentially expressed genes between disease cases and healthy controls using measured transcriptome data. However, the difficulty of measuring gene expression, the limited availability of large samples of disease relevant tissues, and confounding environmental factors all contribute to the mixed results of gene expression studies to date [15]. For BD, for which the most relevant tissue is brain, there is an obvious challenge in collecting large samples for gene expression studies. A recent combined analysis of publically available BD gene expression studies in brain included only 57 unique cases and 60 unique controls. This study found 382 genes to be significantly differentially expressed in brain between BD cases and controls (p-value ≤ 0.05) [15]. While this study has a larger sample size than previous studies, the confounding of gene expression changes as a risk factor for disease versus those that occur as a consequence of disease and drug treatment cannot be removed. This reverse causality issue severely limits the utility of traditional differential expression approaches in dissecting the etiology of this disease.

Recognizing the relevance of gene expression as a contributor to complex disease risk and the difficulties of measuring gene expression directly in disease relevant tissues and in large enough samples, we applied a recently published method, PrediXcan, to identify new genes associated with BD [16]. Using prediction models of expression, we predict the genetic component of gene expression in two large case-control studies for bipolar disorder, and then associate predicted gene expression levels with disease. Among the advantages of the PrediXcan approach are that a) only data on genetic variation and the phenotype are needed; gene expression levels are predicted using genetic data so that it can be applied to any existing GWAS/sequencing study, b) the gene-based results have biological interpretation, c) the directionality of the effects can guide follow-up studies and drug development, d) reverse causality can be for the most part ruled out since disease status and drug treatment do not modify germ line DNA, and e) the multiple testing burden is substantially reduced (from ~1 million to ~10 thousand) [16].

In the current study we develop predictors of gene expression in 10 brain regions and apply these predictors to identify novel genes associated with BD risk using GWAS data from the Wellcome Trust Case-Control Consortium (WTCCC) and Genetics Association Information Network (GAIN). Through this analysis we replicate the association between predicted expression of *PTPRE* and BD risk in whole blood and found a new genome-wide significant association with *BBX* (bobby sox homolog) in the anterior cingulate cortex region of the brain.

## Methods

### Predictors of Gene Expression

For our whole blood analysis, we used gene expression prediction models developed in our original PrediXcan publication [16] based on the Depression Genes and Networks (DGN) transcriptome dataset [17]. This dataset included gene expression information on 922 individuals. These predictors are based on SNPs in the vicinity of each gene (+/‐ 1MB), and used elastic net regularized regression with α = 0.5 [16,18]. We only included common (MAF > 0.05) HapMap Phase 2 SNPs in our predictive models since it was shown that the improvement in cross-validated prediction performance was only marginal when using a more dense set of variants. We restricted analysis to protein-coding genes that had a R^2^ ≥ 0.01 between predicted and measured expression based on 10-fold cross-validation. In total, we tested gene expression models for 8,505 genes based on this dataset (Table 1) [16].

To explore the tissue-context specificity of gene expression changes associated with BD risk, we developed similar predictors of gene expression using the brain region gene expression data from the Genotype-Tissue Expression (GTEx) Consortium (version 6). This included gene expression data from 10 regions of the brain—anterior cingulate cortex (BA24), caudate basal ganglia, cerebellar hemisphere, cerebellum, cortex, frontal cortex (BA9), hippocampus, hypothalamus, nucleus accumbens basal ganglia, and putamen basal ganglia— on 73-103 individuals (varies by region) (Table 1). Consistent with other GTEx analyses, we used normalized measures of expression, reads per kilobase per million (RPKM), and corrected for 15 PEER factors [19], the first 3 genotype-based principal components, and gender [10]. For each brain region, we developed predictors of gene expression using the same method and input parameters as the whole blood gene expression predictors described above [16,18]. We assessed the quality of our gene expression predictors using 10-fold cross validation. To generate gene expression predictors we used the glmnet package implemented in R (R-project.org). As before, we restricted analysis to include only genes with an R^2^ ≥ 0.01 between predicted and measured expression based on the 10-fold cross validation (Table 1).

**Table 1.** Gene Expression Datasets. The gene expression datasets used to generate predictive models of gene expression for PrediXcan analysis. For each tissue, we restricted our PrediXcan analysis to genes that had an R^2^ ≥ 0.01 based on 10-fold cross validation.

| Tissue | Sample Size | Number of Genes | Bonferroni Corrected Significance Threshold |
| --- | --- | --- | --- |
| DGN: Whole Blood | 922 | 8,501 | $5.88 \times 10^{-6}$ |
| GTEXBrain: Anterior Cingulate Cortex (BA24) | 72 | 4,848 | $1.03 \times 10^{-5}$ |
| GTEXBrain: Caudate Basal Ganglia | 100 | 5,137 | $9.73 \times 10^{-6}$ |
| GTEXBrain: Cerebellar Hemisphere | 89 | 5,683 | $8.80 \times 10^{-6}$ |
| GTEXBrain: Cerebellum | 103 | 6,134 | $8.15 \times 10^{-6}$ |
| GTEXBrain: Cortex | 96 | 5,200 | $9.62 \times 10^{-6}$ |
| GTEXBrain: Frontal Cortex (BA9) | 92 | 4,981 | $1.00 \times 10^{-5}$ |
| GTEXBrain: Hippocampus | 81 | 4,697 | $1.06 \times 10^{-5}$ |
| GTEXBrain: Hypothalamus | 81 | 4,595 | $1.09 \times 10^{-5}$ |
| GTEXBrain: Nucleus Accumbens Basal Ganglia | 93 | 5,000 | $1.00 \times 10^{-5}$ |
| GTEXBrain: Putamen Basal Ganglia | 82 | 4,925 | $1.02 \times 10^{-5}$ |
| Total number of gene-tissue pairs | | 59,701 | $8.38 \times 10^{-7}$ |

We also calculated the heritability of gene expression for each gene across the 10 brain regions from GTEx. The heritability of gene expression serves as an upper bound for how well we can predict the trait using only genetic data. We estimated the narrow-sense heritably of gene expression for each gene and each brain tissue using a variance-components model with the genetic relatedness matrix (GRM) estimated from the genetic data, as implemented in GCTA version 1.24.4 [20]. As with our prediction models, we restricted our analysis to include only SNPs in the vicinity of the gene (+/‐ 1MB). As our estimate of the local heritability, we calculated the proportion of variance in gene expression explained by these SNPs using the following mixed effects model:

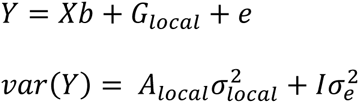

Where *Y* is the gene expression trait, *b* is the vector of fixed effects, and *A*_*local*_ is the GRM calculated from SNPs in the vicinity of each gene. *G*_*local*_ is the combined genetic effect of local SNPs on the gene expression trait with 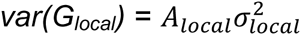. Using this model, the narrow-sense heritability is defined as

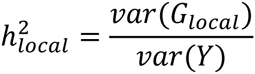

Due to the small sample sizes for our transcriptome datasets in brain, we were unable to estimate the genetic effect of SNPs not in the vicinity of the gene accurately (i.e. a global estimate of h^2^ using genome-wide SNPs).

### GWAS datasets

Our discovery sample included 1,998 BD cases and 3,004 healthy controls of European ancestry genotyped as part of the WTCCC phase 1 study (WTCCC) [21]. Our replication samples included 1,001 BD cases and 1,033 healthy controls of European ancestry genotyped as part of the GAIN study [22]. We used WTCCC genotypes imputed using the 1000 genomes phase 3 reference panel on the UM imputation server [16,23,24]. GAIN genotypes were imputed to include HapMap SNPs [22]. For each dataset, we restricted analysis to include SNPs with high quality imputation (R^2^ ≥ 0.8). Further details about each study can be found in the original publications for each study [21,22].

For each of the datasets, WTCCC and GAIN, we calculated the top 3 principal components based on the imputed genotype data using plink (version 1.90) [25] and GCTA (version 1.24.4) [20]. We included principal components in our analysis to correct for hidden population structure.

### Application of PrediXcan

For each tissue of interest, we estimated the genetically regulated component of gene expression in the WTCCC and GAIN datasets using the gene expression models described above. We then tested for an association between predicted gene expression and BD disease risk in each dataset using logistic regression in R (R-project.org), correcting for the first 3 principal components of the genotype data. We considered any association result genome-wide significant if it had an association p-value ≤ 0.05 after Bonferroni correction for the number of genes tested for the tissue.

### Meta-Analysis

We combined the results of the PrediXcan analysis from our two BD datasets using a fixed effects meta-analysis weighting by sample size in R:

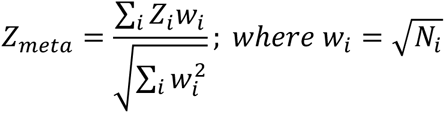

where *Z*_*i*_ is the study specific z-score and *N*_*i*_ is the sample size for each study. We did this separately for each tissue tested. We also checked for a consistent direction of the association effect for each gene between the WTCCC and GAIN datasets.

## Results

### Brain Region Prediction Models

Using data from the GTEx Consortium for 10 brain regions, we generated predictive models of gene expression for 4,599-6,139 protein-coding genes depending on the specific brain region with an R^2^ ≥ 0.01 based on 10-fold cross-validation (Table 1). Given the close to ten-fold difference in sample size for brain regions relative to whole blood (72-103 versus 922), we were not surprised by the smaller number of genes with good predictive models (8,501 in whole blood). In total, we tested 16,173 unique protein-coding genes in at least 1 brain region. On average, each gene was predicted in 3 brain regions.

Despite the smaller sample sizes for each brain region transcriptome dataset, we were able to develop models of gene expression that resulted in significant cross-validated R^2^ between predicted and observed gene expression for a large number of genes (Table 1). Reassuringly, we observed a significant correlation between cross-validated R^2^ and local estimates of h^2^ (Fig 1). The overall correlation between cross-validated R^2^ and local estimates of h^2^ across all genes and tissues was 0.39 (p < 10^−16^). When we further restrict to genes with a local estimates of h^2^ significantly greater than 0 (p < 0.1), the correlation between cross-validated R^2^ and local h^2^ increases to 0.61 (p < 10^−16^). This is consistent with the fact that genes with higher heritability are better predicted. Local h^2^ estimates serve as a benchmark for our ability to predict gene expression traits using our SNP-based models, thus seeing a strong positive relationship between prediction R^2^ and heritability suggests that are models are able to capture the genetic component of gene expression. This relationship between crossvalidated R^2^ from our gene expression models and local h^2^ was seen previously for whole blood [16]. Cross-validated R^2^ is less than the estimated local h^2^ for 39% of genes (points below the red line in Fig 1), suggesting there is room for improvement in our predictive models as the sample sizes for brain expression datasets increase.

**Figure 1.**
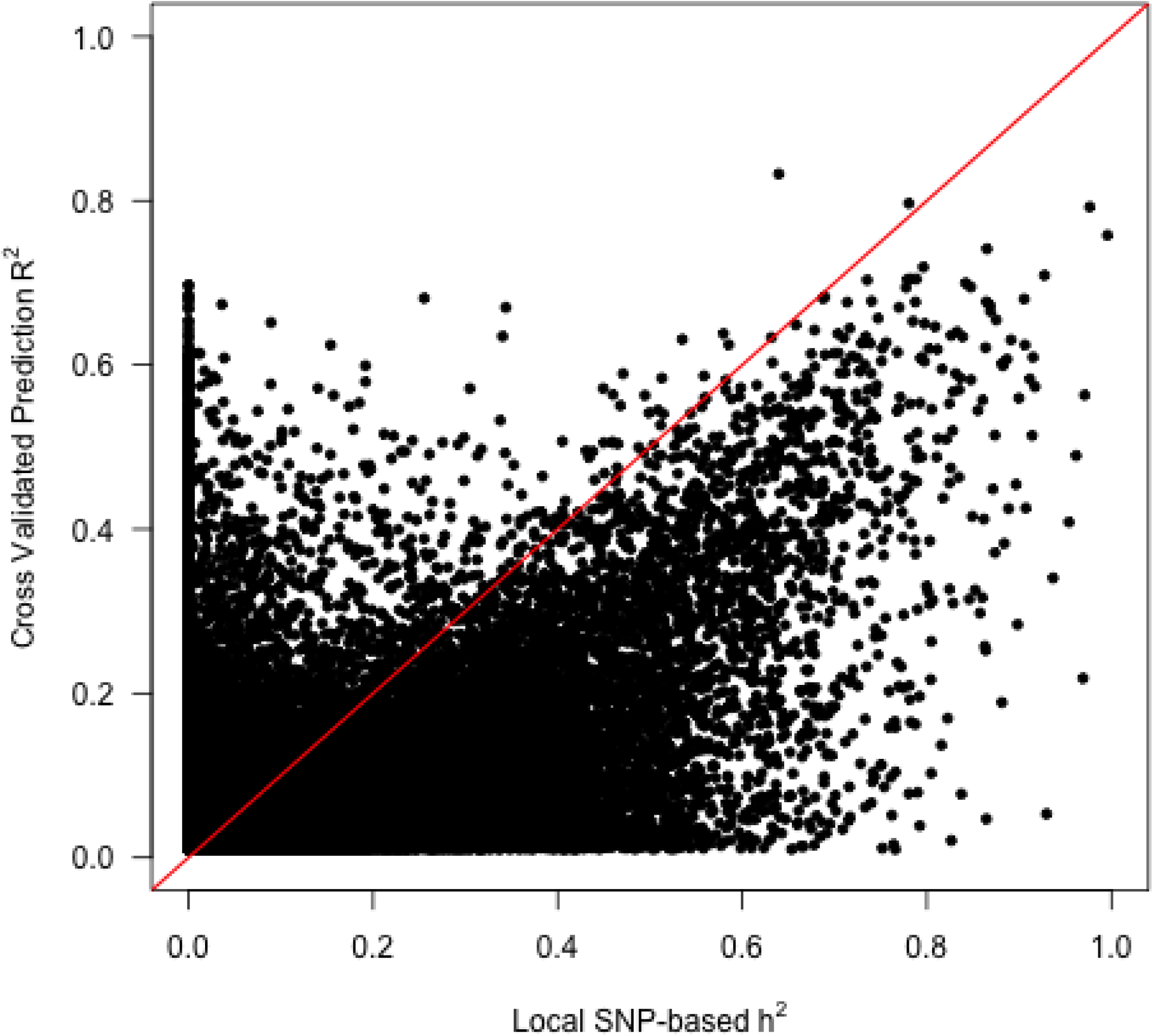
Local Heritability and Prediction Quality. The R^2^ between predicted and measured gene expression based on 10-fold cross-validation for each gene and tissue is plotted against the estimate of local heritability, h^2^. The red line shows R^2^ = h^2^. The local estimate of heritability serves an estimate of the upper bound of our prediction ability for each gene using our SNP-based models of gene expression.

### Association between predicted expression and BD status - Whole Blood

Results from our analysis of the WTCCC sample were previously published along with 6 other disease traits from this dataset [16]. In the WTCCC study we found higher predicted expression of *PTPRE* is associated with increased risk of BD (Table 2, p_WTCCC_ = 8.13 x 10^−7^). In this analysis, we replicated the finding in the GAIN study, which is independent from the WTCCC sample. As in WTCCC, higher predicted expression of *PTPRE* is also associated with BD risk (Table 2, p_GAIN_ = 0.046) in the GAIN dataset. The meta-analysis yields a more significant p-value than in the discovery set, strengthening the evidence for our results (Fig 2 and Table 2, p_meta_ = 1.92 x 10^−7^). A gene-level Manhattan plot of the combined PrediXcan results across multiple tissues including whole blood is presented in Fig 2, whereas the whole blood specific results are in Sup Fig 1A and Sup Fig 2A. The association between *PTPRE* and BD risk remains significant after correcting for the total number of gene-tissue pairs tested. We note that the genome‐ and tissue-wide significance threshold when correcting for all gene tissue pairs tested is 8.37 x 10^−7^ (0.05/total number of gene/tissue combinations; blue line in Fig 2) while the single tissue genome-wide significant threshold for whole blood is 5.88 x 10^−6^ (0.05/# genes in the tissue; Table 1). Notice that in general, our significance thresholds are conservative because we do not account for the correlations between expression values across tissues and genes, which would reduce the effective number of independent tests. *PTPRE* encodes the protein tyrosine phosphatase, receptor type E and is involved in RAS signaling, SATA signaling, and activation of voltage-gated K+ channels [26,27]. Interestingly, an intronic SNP in *PTPRE* was previously found to be suggestively associated with response to the stimulant amphetamine [28,29]. However, the reported SNP, rs12049671, is not an eQTL for *PTPRE* in whole blood (p-value > 0.05).

**Table 2.** Top PrediXcan Results. Genome-wide significant genes in the meta-analysis of WTCCC and GAIN after correcting for multiple testing within each tissue.

| Gene | Tissue | WTCCC |  | GAIN |  | Meta-Analysis p-value |
| --- | --- | --- | --- | --- | --- | --- |
|  |  | z-stat | p-value | z-stat | p-value |  |
| *PTPRE* * | DGN: Whole Blood | 4.93 | $8.13 \times 10^{-7}$ | 1.99 | 0.047 | $1.92 \times 10^{-7}$ |
| *BBX* * | GTEXBrain:<br>Anterior Cingulate<br>Cortex (BA24) | -4.49 | $7.02 \times 10^{-6}$ | -2.83 | $4.68 \times 10^{-3}$ | $1.11 \times 10^{-7}$ |
| *BBS4* | GTEX Brain:<br>Putamen Basal<br>Ganglia | 3.29 | $1.02 \times 10^{-3}$ | 3.15 | $1.66 \times 10^{-3}$ | $7.55 \times 10^{-6}$ |

\*Meta-analysis p-values remain significant after correcting for the total number of tests conducted by gene and tissue

**Figure 2.**
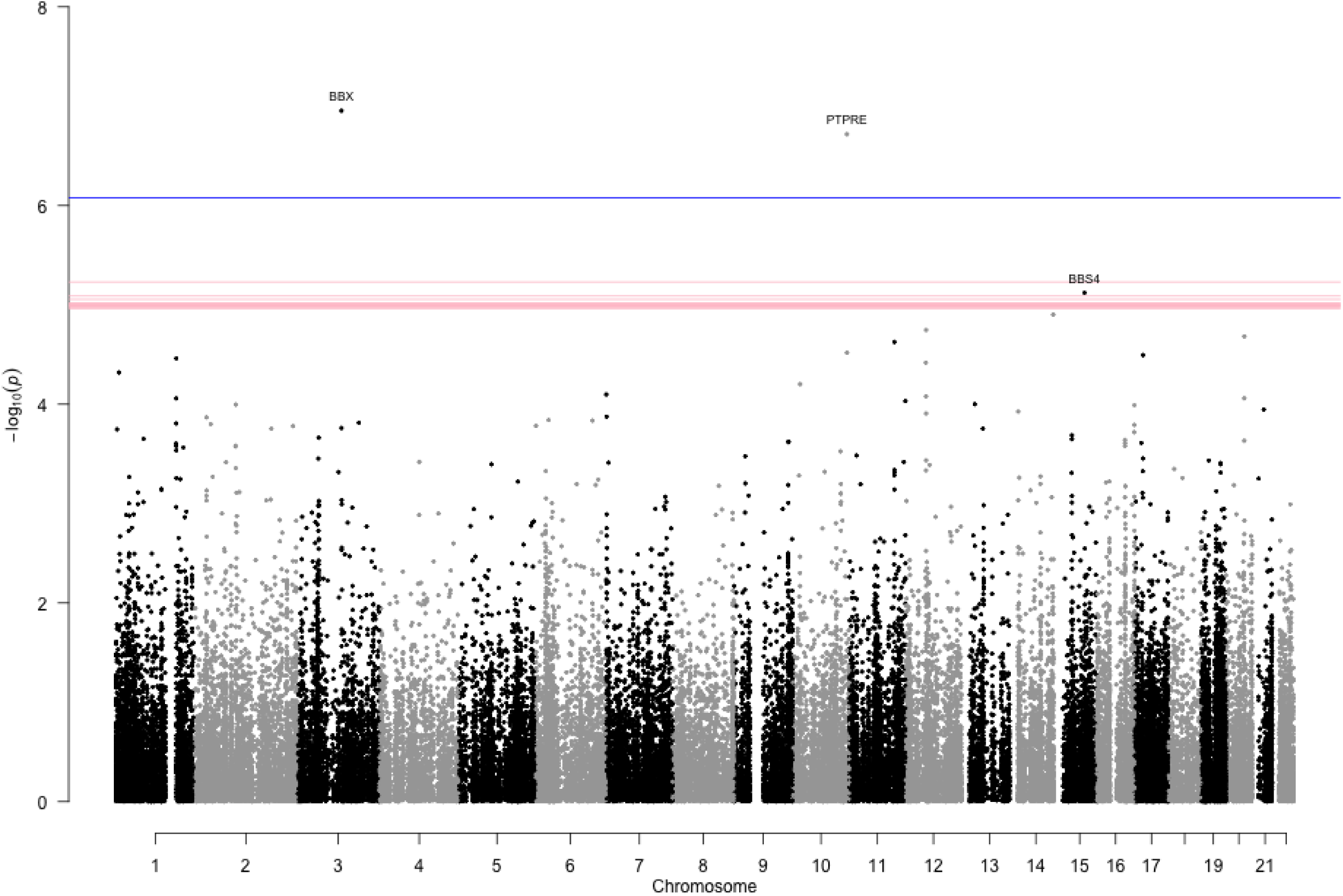
PrediXcan Results Across Tissues. The p-values plotted by genomic position for all of the PrediXcan meta-analysis results across 11 tissues (whole blood and 10 brain regions). The blue horizontal line represents the Bonferroni corrected significance threshold after correcting for all genes and all tissues. The red lines are the tissue-specific Bonferroni corrected significance thresholds. The top genes that remain significant after correcting for multiple testing within each tissue are labeled.

### Association between predicted expression and BD status - Brain Regions

In our PrediXcan analysis in brain we identified 4 genes associated with BD risk in the WTCCC sample after correcting for multiple testing‐‐*BBX* in anterior cingulate cortex (BA24), *KIF1A* in cerebellar hemisphere, *TDRD9* in cortex, and *ITGA4* in caudate basal ganglia. Of these, we were able to replicate the association between *BBX* and BD risk in the anterior cingulate cortex (BA24) using the GAIN dataset. The meta-analysis results for individual brain regions are presented in Sup Figs 1B-K and Sub Figs 2B-K. We found that lower predicted expression of *BBX* is associated with increased risk of BD after Bonferroni correction for the number of genes and tissues tested (Table 2, p_WTCCC_ = 7.02 x 10^−6^; p_GAIN_ = 0.0047; p_meta_ = 1.11 x 10^−7^). *BBX*, bobby sox homolog, is an HMG box containing transcription factor important for cell cycle progression. While not genome-wide significant, an intronic variant in *BBX*, rs6437740 was reported to be suggestively associated with smoking behavior [30].

Our meta-analysis of the WTCCC and GAIN datasets resulted in one additional gene reaching genome-wide significance, but did not pass our more stringent threshold correcting for the number of tissues as well. Higher predicted expression of *BBS4* showed a suggestive association with increased BD risk in the putamen basal ganglia (Fig 2 and Table 2, p_WTCCC_ = 0.0010; p_GAIN_ = 0.0016; p_meta_ = 7.55 x 10^−6^). While this association still needs to be replicated in an independent dataset, *BBS4* represents an interesting candidate gene for BD risk. Recessive null mutations in *BBS4*, Bardet-Biedl Syndrome 4, are known to cause Bardet-Beidle Syndrome, a disease characterized by retinitis pigmentosia, obesity, kidney dysfunction, polydactyly, behavioral dysfunction, and hypogonadism (OMIM: 615982). Furthermore, members of the BBS gene family are involved in proper cilia formation and function.

We provide the full set of our PrediXcan results for whole blood and the 10 brain regions in Supplemental Table 1.

## Discussion

PrediXcan is a recently published gene-level association method that tests the mediating effects of gene expression levels in influencing phenotype. Unlike standard gene mapping approaches, PrediXcan interrogates genes, not single variants, and tests a specific biological mechanism ‐‐ that altered gene expression is associated with disease risk. We applied this approach to identify genes associated with BD risk using predicted gene expression in whole blood and 10 different brain regions. Despite the smaller sample sizes in our expression datasets for brain tissues, we were able to achieve high quality predictive models for many genes (Table 1). Thus, we were able to apply PrediXcan to this subset of genes.

Our PrediXcan analysis identified and replicated 2 new genes associated with BD risk. We found a reproducible association between higher *PTPRE* expression in whole blood and increased risk of BD (Table 2). *PTPRE* is known to be involved with activation of the RAS signaling pathway. Disruption of this pathway has already been associated with psychiatric diseases including autism spectrum disorders (ASD). Other genes in the RAS signaling pathway have been associated with psychiatric diseases-*CACNA1C, SYNGAP1*, and *MAPK3* to name a few [31,32]. Interestingly, while hyper activation of RAS signaling appears to be a risk factor for ASD, there is some evidence for hypo activation of RAS signaling in a subset of BD cases with sleep disturbances [32]. Thus, increased levels of *PTPRE*, an inhibitor of RAS signaling, as a risk factor for BD is consistent with prior knowledge about the relationship between psychiatric disease and the RAS signaling pathway.

In the anterior cingular cortex (BA24) region of the brain, we found that lower *BBX* expression is associated with increased BD risk (Table 2). Interestingly, this region of the brain is thought to be involved in a variety of autonomic and higher-level functions like reward anticipation, decision-making, empathy, impulse-control, and emotion [33]. Furthermore, individuals with damage to this region often display psychopathic and sociopathic behaviors [33]. Thus, this region of the brain is likely relevant to psychiatric disease risk. *BBX* is a member of the high motility group (HMG)-box family of transcription factors. Previous zebrafish and mouse studies showed that expression of BBX was focused in the CNS and brain, and that the gene played a role in neurodevelopment [34].

Finally, we found an additional (suggestive) association between predicted expression of *BBS4* and risk of BD in the putamen basal ganglia (Table 2). The association between *BBS4* and BD risk does not remain significant after correcting for all genes and tissues tested however we note that our Bonferroni corrected significance threshold is conservative because it does not account for correlation in gene expression (which would reduce the effective number of independent tests). Mutations in *BBS4* are known to cause Bardet-Biedl Syndrome 4. Using data from millions of medical records, Blair *et al*. found a number of complex diseases were comorbid with Bardet-Biedl Syndrome 4, including bipolar disorder, schizophrenia, obsessive compulsive disorder, intellectual disability, epilepsy, and sleep disorder [35]. While our observed association between *BBS4* expression and BD risk needs to be replicated, based on what we know about the Mendelian disease, *BBS4* is an excellent candidate gene for BD and general psychiatric disease risk.

In our overall PrediXcan analysis using the WTCCC data, we note a small degree of inflation in our p-value distribution that impacted the results from our metaanalysis (Sup Fig 2). The PrediXcan results from the GAIN dataset did not show a similar inflation. This inflation could be due to a small degree of population structure or differences between cases and controls in the WTCCC dataset. Because PrediXcan is based on combining information from SNPs that are associated with gene expression, another source of inflation could be due to the fact that this set of SNPs is enriched for true disease associations similar to the observations seen with eQTLs [13,14]. Reassuringly, even after adjustment for this inflation using genomic control, our results remain genome-wide significant.

The PrediXcan approach has many advantages and limitations compared to current gene mapping strategies. Unlike GWAS, where the biological interpretation of results is a major challenge, PrediXcan tests a specific mechanistic hypothesis about the influence of altered gene expression on disease risk. Thus, the results are immediately interpretable and can be readily tested in cell-based and model organism follow-up studies. In contrast to differential gene expression experiments, PrediXcan method is not plagued with reverse causality issues.

The performance of our method was limited by the quality of the prediction models. Our comparison with estimates of heritability shows that as sample sizes grow in GTEx and other brain tissue expression datasets, we will be able to improve our ability to identify brain specific genes.

In contrast to widespread prior belief, recent publication from the GTEx consortium demonstrated that there is a large component of genetic regulation that is shared across multiple tissues [10]. Consequently, genes identified in whole blood and other more readily available tissues are still relevant to brain related disease phenotypes. For example, the association between *PTPRE* and BD risk identified using whole blood models recapitulate when brain tissues are used with equal direction of the effects albeit with different degree of significance likely due to the limited sample sizes used in training the models (Table 3). It is important to note that the predictors developed in whole blood used an independent training transcriptome dataset than the predictors for brain (DGN vs. GTEx).

**Table 3.** PTPRE PrediXcan Results. Results for the association between predicted expression of *PTPRE* and BD risk for the tissues with sufficient data. Only 4/10 of the available brain regions had quality predictors for *PTPRE* and are presented below. While less significant, the brain tissues show a consistent direction of effect as whole blood.

| Tissue | WTCCC |  | GAIN |  | Meta-Analysis<br>p-value |
| --- | --- | --- | --- | --- | --- |
|  | z-stat | p-value | z-stat | p-value |  |
| DGN: Whole Blood | 4.93 | $8.13 \times 10^{-7}$ | 1.97 | 0.047 | $1.92 \times 10^{-7}$ |
| GTEXBrain: Frontal Cortex (BA9) | 3.66 | $2.52 \times 10^{-4}$ | 2.03 | 0.042 | $3.03 \times 10^{-5}$ |
| GTEXBrain: Cortex | 2.59 | $9.60 \times 10^{-3}$ | 0.81 | 0.42 | $9.23 \times 10^{-3}$ |
| GTEXBrain: Cerebellum | 2.58 | $9.98 \times 10^{-3}$ | 0.043 | 0.97 | 0.030 |
| GTEXBrain: Anterior Cingulate Cortex (BA24) | 1.36 | 0.175 | 1.77 | 0.077 | 0.035 |

By applying PrediXcan to existing GWAS datasets we identified novel BD risk genes and propose the potential mechanism underlying these associations. Our results further highlight the relevance of gene expression to complex disease risk [13,14,36]. Furthermore, for BD, we add to the existing understanding of disease biology and provide support for the importance of the RAS signaling pathway for psychiatric disease [31,32,37].

## Acknowledgements

We acknowledge the following grants that supported this work: T32 MH020065 (K.P.S.), P50 MH094267 (Conte), R01MH107666 (HKI), K12CA139160 (HKI), P30DK020595 (DRTC), R01MH101820 (GTEx). The datasets used for the analyses described in this manuscript were obtained from dbGap at http://www.ncbi.nlm/nih.gov/gap through dbGaP accession numbers phs000424.v6.p1 (GTEx) and phs000017.v3.p1.c1 (GAIN). This study makes use of data generated by the Wellcome Trust Case-Control Consortium (WTCCC). A full list of investigators who contributed to the generation of this data is available from www.wtccc.org.uk. Funding for the project was provided by the Wellcome Trust under award 076113 and 085475.

---

**Supplemental Figure 1. PrediXcan Results by Tissue.** The p-values plotted by genomic position for the PrediXcan meta-analysis results for whole blood (A) and the 10 brain regions (B-K). The red line in each plot is the tissue-specific Bonferroni corrected significance threshold. Genes that remain significant after correcting for multiple testing within each tissue are labeled.

**Supplemental Figure 2. QQ-plots for PrediXcan Results by Tissue.** QQ-plots showing the p-value distribution for each of our PrediXcan meta-analyses by tissue for whole blood (A) and the 10 brain regions (B-K). Each plot shows the 95% confidence interval under the null distribution in gray and the thresholds for various false-discovery rates (dashed lines). The blue horizontal line represents the tissue-wide Bonferroni corrected significance threshold.

**Supplemental Table 1. PrediXcan results by tissue.** Results for all genes across all tissues tested for association with BD risk.

